# What Strengthens Protein-Protein Interactions: Analysis and Applications of Residue Correlation Networks

**DOI:** 10.1101/2023.03.15.532709

**Authors:** Ta I Hung, Yun-Jung Hsieh, Wei-Lin Lu, Kuen-Phon Wu, Chia-en A. Chang

## Abstract

Identifying critical residues in protein-protein binding and efficiently designing stable and specific protein binders is challenging. In addition to direct contacts in a protein-protein binding interface, our study employs computation modeling to reveal the essential network of residue interaction and dihedral angle correlation critical in protein-protein recognition. We propose that mutating residues regions exhibited highly correlated motions within the interaction network can efficiently optimize protein-protein interactions to create tight and selective protein binders. We validated our strategy using ubiquitin (Ub) and MERS coronaviral papain-like protease (PLpro) complexes, where Ub is one central player in many cellular functions and PLpro is an antiviral drug target. Our designed UbV with 3 mutated residues resulted in a ∼3,500-fold increase in functional inhibition, compared with the wild-type Ub. Further optimization by incorporating 2 more residues within the network, the 5-point mutant achieved a K_D_ of 1.5 nM and IC_50_ of 9.7 nM. The modification led to a 27,500-fold and 5,500-fold enhancements in affinity and potency, respectively, as well as improved selectivity, without destabilizing the UbV structure. Our study highlights residue correlation and interaction networks in protein-protein interaction, introduces an effective approach to design high affinity protein binders for cell biology and future therapeutics solutions.

## Introduction

Understanding what drives protein-protein binding and efficiently selection of which residues of a protein to modify to increase protein–protein interactions (PPIs) are the key to design a protein binder targeting its binding protein ^1,2^. Strategies that can efficiently and accurately identify residues for enhancing PPIs also have broad applications in therapeutics and cell-biology studies. Knowledge-based, physics-based, and data-driven methods have been developed for investigating protein-protein interactions and selection of mutations to enhance PPIs ^3–9^. Combined computational and combinatorial libraries or *in vitro* evolution approaches are also popular strategies in protein engineering to design stable and specific protein binders ^10–12^. Importantly, utilizing a handful of mutated residues to achieve significantly enhanced PPIs lowers the possibility of introducing unstable engineered proteins. However, the task is even more challenging because it requires highly integrated molecular modeling and experimental techniques to understand PPIs for re-engineering a protein to increase the protein-protein binding affinity.

PPI networks are extremely complicated, and selecting a good target system for modifications requires specialized expertise. Here we chose Ubiquitin (Ub) as our target system, which plays critical roles in numerous biological functions ^13^. Ub is a 76-residue small protein associated with post-translational modifications. This regulatory protein canonically binds to its cascade E1-E2-E3 enzymes for ubiquitination and Ub chain formation to modify nearly half of the human proteome ^14,15^. Conversely, deubiquitinase (DUB) ^16^ cleaves the covalent isopeptide bonds from the Ub chains or substrates to release Ub and substrates. The interactomics of Ub and cellular proteins have been intensively accessed ^15^ revealing that accurate regulations of Ub networks govern the cellular fates.

Misregulation of these enzymes substantially affects cellular functions, hence leading to diseases such as cancers ^17^. Moreover, DUBs have been identified in several viral genomes as a tool interfering in the host antiviral defense. For example, Papain-like protease (PLpro) of coronavirus (CoV) is classified as a viral DUB specific to Ub and the Ub-like ISG15 ^18,19^. Studies show that PLpro alters the innate immune responses, which contributes to the rapid spread of CoVs (MERS, SARS-CoV, SARS-CoV-2) ^20–24^, thus causing pandemic deaths and decreasing the global economy ^25^.

PLpro of MERS and SARS-CoV-2 are crucial for viral replication via proteolytic cleavage of viral nonstructural proteins (NSPs). The PLpro domain resides in NSP3, which drives viral genome replication and subgenomic RNA synthesis ^26,27^. PLpro can recognize and cleave the NSP1-2, NSP2-3 and NSP3-4 junctions after the amino acid sequence LXGG to yield functional viral proteins as well as perform deubiquitination and deISGylation ^23,24,28^. Deubiquitination and deISGylation alter host signaling pathways critical for inducing cellular antiviral and pro-inflammatory innate immune responses and eventually suppresses the antiviral response ^28,29^. Therefore, inhibition of PLpro simultaneously disrupts the viral replication and prevents PLpro from disrupting the innate immune response. With both properties, PLpro is an ideal antiviral drug target.

Importantly, wtUb exhibits high thermostability (T_m_ > 90°C) and so is a great template for protein design. The re-engineered Ub also has potential advantages such as good binding specificity to PLpro and easy synthesis as compared with chemical compounds. Phage-display screened Ub variants (UbVs) against cognate enzymes including MERS PLpro demonstrated their feasibility in regulating the activities of E3 ligases and DUBs ^30–32^. The phage-display screening technique focused on three surface patches of Ub to iteratively mutate and select tight binders. The raised DUB UbVs are great inhibitors exhibiting IC_50_ at a range of 1-30 nM ^31–33^. Alternatively, computational results were used to rationally design a screening library for identifying tight binding regulator UbVs for USP7 ^34^ and USP21^35^. A combined computational and phage display screening of UbVs targeting USP7 resulted in K_D_ of variant U7Ub25.2540 at 56 nM whereas the K_D_ for wtUb-USP7 is greater than 200 μM. A pool of designed 6,000 UbVs for USP21 reveals approximately 10% of the variants are tight binders for USP21 consistently between experimental and computational screenings. However, in silico screening of such large pairing UbV-USP21 requires intensive computational resources. The expensive and time-consuming empirical screening hampers design of protein-based inhibitors.

Here we present an integrated computational and experimental approach to identify critical regions in protein-protein binding, which have highly correlated motions and the regions are connected by an interaction network. We demonstrated that mutating residues in these regions can efficiently optimize protein-protein interactions to create tight and selective protein binders. We showed that our designed UbVs with 2 to 3 mutated residues achieved 3,500-fold inhibitory efficiency and binding affinity as compared with the wtUb to MERS PLpro (Table 1). MERS PLpro cleaves both K48 and K63-linked Ub chains ^18,23^, and it has distinct inhibitor recognition specificity different from SARS-CoV and SARS-CoV-2 ^36^. We used non-covalent amino acid interaction networks of the Ub and MERS PLpro (Ub-PLpro) complex to guide the design of UbVs for increasing the UbV–PLpro binding affinity that inhibits the function of PLpro. The initial design began with 2-point mutations for cost-efficient experiments and for keeping an intact overall complex structure. Binding affinity K_D_ and IC_50_ measurement confirmed the inhibition of our designed dual-mutation UbVs and suggested additional UbVs. Integrating both experimental data and computation analysis re-informed the design to yield more UbVs for experiments (Fig. 1).

**Table 1.** Computational and experimental evaluation of binding affinity between MERS-CoV-PLpro and UbVs.

| Ubiquitin and variants | Binding Energy (kcal/mol) | Dissociation Time (ns) | IC<sub>50</sub> | K<sub>D</sub> | PLpro selectivity* |  |
| --- | --- | --- | --- | --- | --- | --- |
|  |  |  |  |  | MERS-CoV | SARS-CoV-2 |
| wtUb | -41.95 ± 2.56 | 32 | 52.91 ± 6.98 μM | 40.75 ± 3.82 μM | - | - |
| A46F | -48.25 ± 1.09 | 67.5 | 1.64 ± 0.03 μM | 2.74 ± 0.29 μM | +++ | - |
| K48E | -49.52 ± 2.56 | 75 | 3.94 ± 0.42 μM | 5.15 ± 0.71 μM | ++ | - |
| E64Y | -42.00 ± 1.49 | 68 | 0.29 ± 0.03 μM | 0.46 ± 0.04 μM | ++++ | - |
| A46F-K48E (**UbV2**) | -50.54 ± 2.35 | 207 | 0.20 ± 0.00 μM | 0.22 ± 0.03 μM | +++++ | - |
| A46F-K48L | -49.03 ± 2.33 | 200 | 0.23 ± 0.01 μM | N/A | N/A | N/A |
| A46F-K48S | -47.13 ± 2.84 | N/A | 0.18 ± 0.01 μM | N/A | N/A | N/A |
| A46F-K48E-G75R | -45.24 ± 1.48 | 40 | 9.74 ± 0.15 μM | 11.41 ± 1.26 μM | N/A | N/A |
| A46F-K48I | N/A | N/A | 0.49 ± 0.04 μM | N/A | N/A | N/A |
| A46F-K48E-E64Y (**UbV3**) | -53.77 ± 1.37 | >250 | 14.84 ± 1.44 nM | 2.77 nM | +++++ | - |
| A46F-K48E-R74N-G75S (**UbV4**) | -50.18 ± 0.88 | 65 | 0.11 ± 0.01 μM | 0.15 ± 0.02 μM | +++++ | - |
| A46F-K48L-R74N-G75S | -46.49 ± 0.98 | >250 | 0.13 ± 0.01 μM | 0.14 ± 0.02 μM | N/A | N/A |
| A46F-K48E-E64Y-R74N-G75S (**UbV5**) | -50.81 ± 3.25 | 88 | 9.71 ± 0.74 nM | 1.48 nM | +++++ | - |
| ME.2** | -47.98 ± 1.72 | >250 | 15.62 ± 2.54 nM | 53.2 ± 2.2 nM | +++++ | - |
| ME.4** | N/A | N/A | 28.57 ± 1.94 nM | 35.9 ± 1.6 nM | +++++ | - |

\* One “+” indicates 0-20% reduction of PLpro activity compared with UbV-free conditions.
\*\* ME.2 and ME.4 were previous UbVs <sup>33</sup> introducing 15 mutation sites independently.
The K<sub>D</sub> values and PLpro selectivity for ME.2 and ME.4 UbVs were extracted from published work <sup>34</sup>.

**Figure 1.**
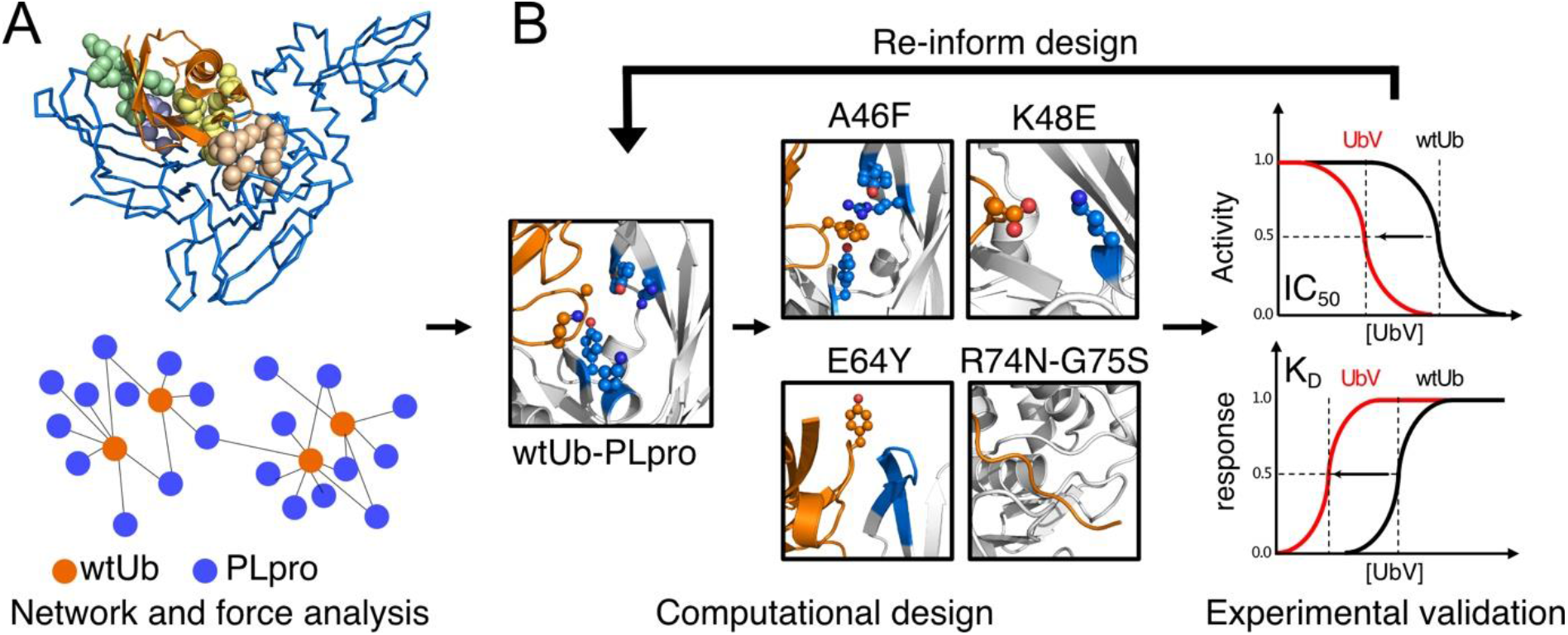
Workflow of rational design of ubiquitin (Ub) variants. **(A)** Molecular modeling with molecular dynamics (MD) simulations and post-analysis to reveal highly correlated regions between the interface of PLpro (presented marine trace) and Ub (orange). **(B)** Integrated computational and experimental design, validation, and interpretation.

## Results and Discussion

### Identification of key residues in the PPI interface

We first performed several 500-ns atomistic molecular dynamic (MD) simulations to model full protein flexibility for the wtUb–PLpro complex. By using combined side-chain dihedral correlation (Fig. S1A) and force distribution analyses (FDA)^37,38^ (Fig. S2), we identified 4 highly correlated regions at the contact interfaces of Ub–PLpro: the hydrophobic core, alpha helix, C-terminal tail of Ub (named Ub-tail), and Zn binding region (Fig. 2). Next, we suggested highly interactive residues with strong correlation in the residue network and also on the Ub–PLpro interface as potential mutation residues in each region (Fig. 2 and see method in Fig. S1). Notably, existing studies showed that altering residues near the C-terminus of Ub hampered its biological function ^39–41^; therefore, we hypothesized that mutating our suggested residues in the Ub-tail could prevent the substrate from ubiquitination. MERS PLpro favors binding and cleaving K48 and K63-linked Ub chains ^18,23^, which implies that residues near the conjugation points interact with PLpro frequently. Of note, K48 and K63 are in the hydrophobic core and Zn binding region, respectively. Mutating residues in the dense interactive spots may easily hamper PPI, which also offers potential in strengthening their interactions. Because the Zn binding and Ub-tail regions are more flexible, we began with modifying A46 and K48 in the hydrophobic core and V70 and R42 in the helix region. We considered side-chain proximity between a UbV and PLpro to utilize unused space or enhance charge-charge attraction to design mutations. Coincidentally, these mutated residues were also reported in existing UbVs ^33^.

**Figure 2.**
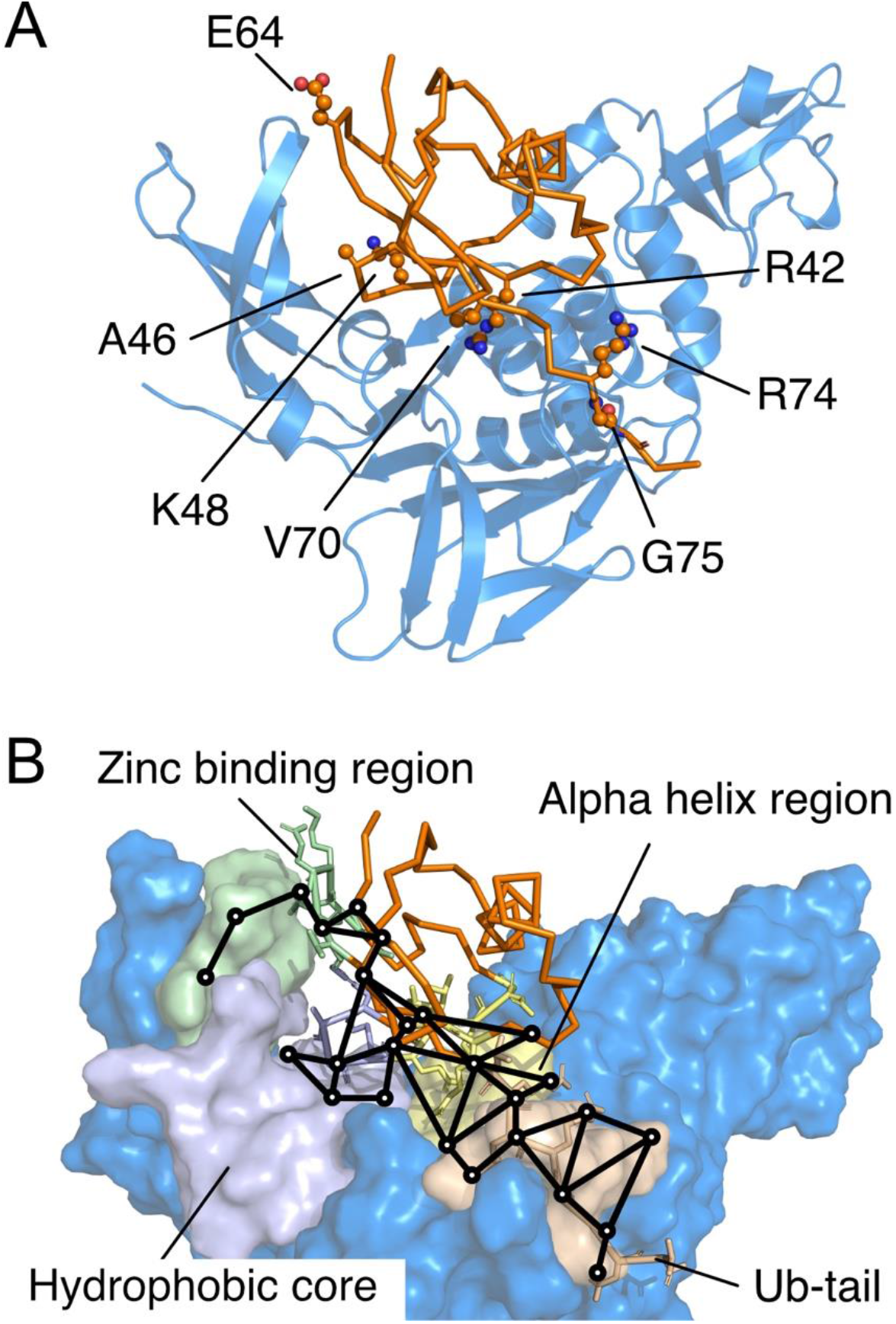
Highly correlated regions between Ub and MERS-CoV PLpro. **(A)** Suggested potential mutation residues based on interaction networks: R42, A46, K48, E64, V70, R74, and G75. Color-coded regions: Ubiquitin (Ub, orange), PLpro (marine), Zn binding region (palegreen), Hydrophobic core (lightblue), Alpha helix region (paleyellow) and Ub-tail (wheat). Note that R42 and V70 were not considered in further Ub variant (UbV) designs and experiments after computational prediction. **(B)** Computed dihedral correlation network showing how mutation leads to conformation changes at distal regions. (See Fig S1A for details)

To predict the intermolecular attractions between each UbV-PLpro, we used molecular mechanics Poisson-Boltzmann surface area (MM/PBSA) and local structural analysis to evaluate PPI energy (Fig. S3) and investigate the local attraction (Fig. S4), respectively. Variants with A46 and K48 mutation led to better UbV–PLpro attractions. However, the K48E-V70E variant could not yield good van der Waal (vdW) interactions and modifying R42 to the oppositely charged residue R42D yielded poor predicted UbV–PLpro interaction energies (Table S1 and Fig. S4). The results suggested that mutating R42 or V70 may not lead to tight binding events. Hence, we first focused on the variants altering A46 and K48. Additionally, for every designed UbV with predicted stronger UbV–PLpro attractions than that of wtUb (Table S1), we also performed PPI Gaussian-accelerated MD (PPI-GaMD) in an explicit solvent model to examine their binding residence time, with protein flexibility and solvent effects also considered ^42^ (Fig. S5). Because PPI-GaMD applied boost potential to enhance conformational sampling, the residence time cannot be compared directly with the binding affinity or dissociation rate constant (*k*_*off*_). Nevertheless, the dissociation time for these UbVs was all longer than that of the wtUb, which suggests that these variants should serve as tighter binders than wtUb.

### Experimental characterizations of inhibitory UbVs

Next, we evaluated the UbV-dependent inhibitory efficiencies for MERS PLpro by using sensitive fluorescent polarization (FP) to monitor the dynamics of fluorescein conjugated to the C-terminus of ISG15 (Fig. 3A). We examined the designed 4 UbVs with dual mutations in positions A46F and K48 (E, S, I and L) as well as 2 single-mutation UbVs, A46F and K48E. The designed UbVs all bound tightly to PLpro (K_D_ in Table 1 and Fig S8) and efficiently inhibited the enzyme function with reduced IC_50_ (Fig. 3B, Table 1 and Fig S7). In comparison to wtUb, substitution of A46 in the hydrophobic core by a non-polar residue PHE successfully formed stronger vdW attractions with nearby residues of PLpro (Fig. 5A and Table S1). For example, A46F induced local conformational arrangements and recruited Y208 and Y223 in PLpro to form π-π interactions and R233 in PLpro to form π-cation attractions (Fig 5A). The A46F mutant substantially stabilizes the interactions of Ub-PLpro, improving K_D_ by 15-fold compared with wtUb (Table S1). The polarity of K48 in wtUb significantly affected both the local interactions and network correlation due to the large repulsive force to the surrounding K204 of PLpro. Mutation of K48 to nonpolar or negatively charged residues such as LEU or GLU should increase the attractive forces with the K204 (Fig. 5A). Notably, the single-point mutations A46F and K48E could achieve an IC_50_ 1.6 and 3.9 μM, respectively, as compared with wtUb (52.9 μM). The 46-48 dual mutants further elevated the inhibition to IC_50_ ∼0.2 μM (Table 1), approximately 250-fold as compared with wtUb. We found a synergistic inhibitory effect and binding affinity by A46F and K48E (or K48L/K48S/K48I) toward the interactions of PLpro and UbV (Fig. 3C, 4A and Fig. S8).

**Figure 3.**
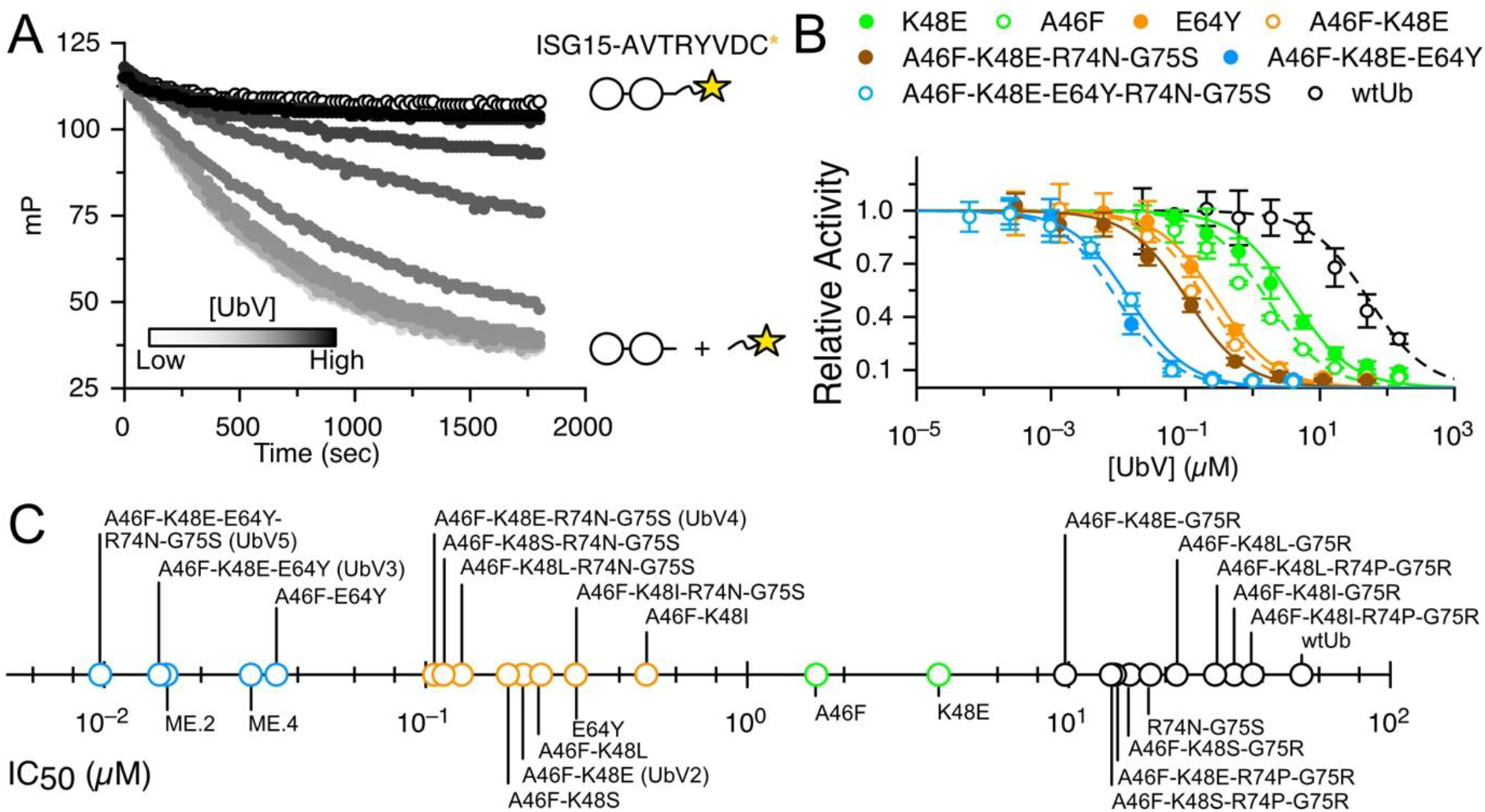
The inhibitory IC_50_ of UbVs. The scheme in (**A**) presents the cleavage reaction of substrate ISG15* where * stands for AVTRYVDC sequence (part of SARS-CoV-2 NSP2) crosslinked with a fluorescent probe fluorescein (star sign). When UbV is added in the mixture, the substrate binding site is blocked and cleavage is retarded. (**B**) The real-time cleavage reaction is monitored by detecting the fluorescence polarization. The black-gray gradient representative relationship of strong-weak inhibitions. IC_50_ curves of selected 7 UbVs and wild-type Ub (wtUb) show PLpro activity progressively inhibited from 2-to 5-point mutations. (**C**) Experimentally measured IC_50_ of 26 UbVs are summarized in a logarithmic scale, with black, green, orange and blue circles indicating weak, medium, strong and very strong inhibitors, respectively.

**Figure 4.**
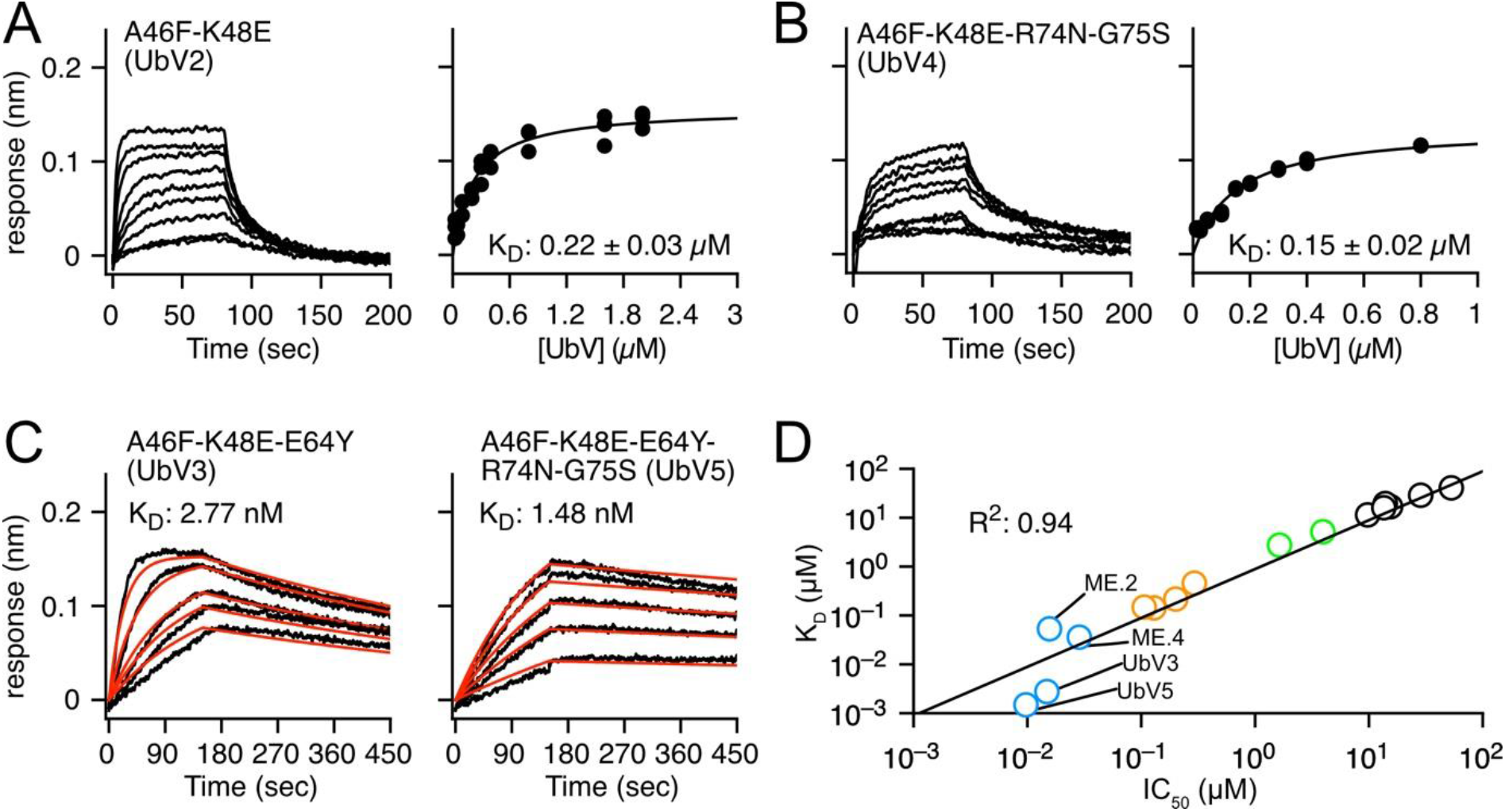
**(A)** BLI sensorgram and fitting curve of UbV2 and UbV4 show K_D_ of 0.22 μM and 0.15 μM, respectively. (**B**) UbV3 and UbV5 bind to PLpro tightly showing K_D_ of 2.77 and 1.48 nM, respectively. The fitting values for k_on_ and k_off_ were colored in red in the titrated BLI sensorgrams **(C)** The measured IC_50_ and K_D_ values are greatly correlated whereas R^2^= 0.94. K_D_ values of ME.2 and ME.4 were published previously^33^.

**Figure 5.**
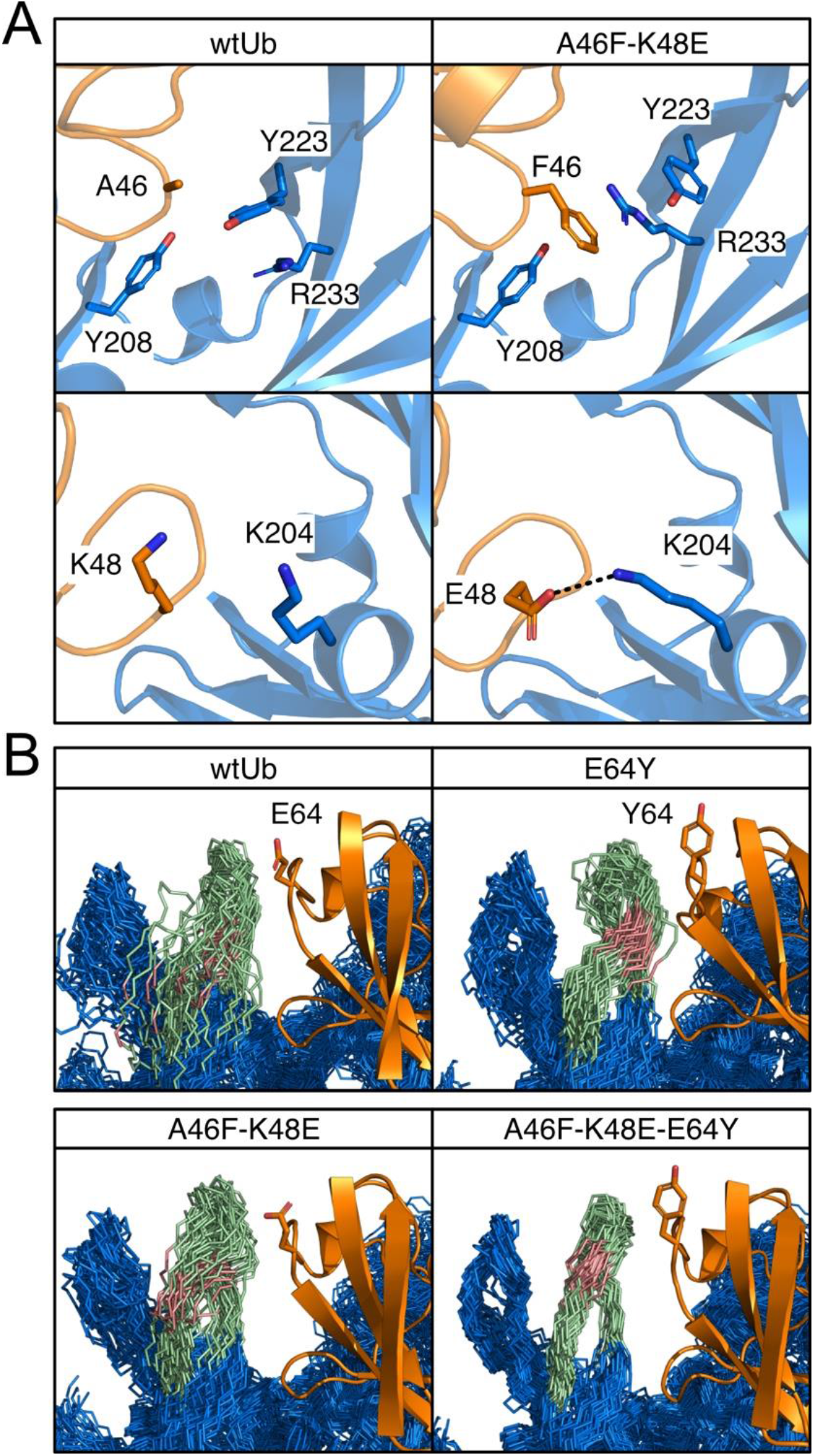
Analysis of conformational dynamics and interactions of wtUb and UbVs for the hydrophobic core and Zn binding region. Ub and PLpro are orange and marine, respectively. **(A)** New attractions introduced by A46F and K48E mutations in the hydrophobic core. **(B)** Superposition of 50 conformations from a 500-ns MD run. The Zn binding region of PLpro (pale green) is highly stabilized by E64Y mutation, where the hinge motion of backbone E230 (salmon) in the interaction network contributes significantly to the loop conformations. Wile-type E64 or mutated Y64 is shown in stick.

We demonstrated that mutating two residues in the hydrophobic core is an effective strategy for inhibitor design, so we further added more mutated residues from our suggested list (Fig. 2). We chose A46F-K48E (termed UbV2), which exhibits good IC_50_ and K_D_, and added one mutated residue, E64Y, from the Zn binding region. Because MERS PLpro explicitly recognizes K63-linked Ub chains, and our residue network shows a strong correlation around S65 (Fig. S1), we conjectured that modifying the nearby residue E64 may contribute to both binding strength and specificity. Experimental measurements for A46F-K48E-E64Y (termed UbV3) showed remarkable binding specificity and inhibition to PLpro, with an IC_50_ of 15 nM, significantly agreeing with the strongest computed binding energy (Table 1). Both the root mean square fluctuation (RMSF) and dihedral entropies of the protein (Fig. S6) were reduced as compared with that of UbV2, so the Zn binding region was largely stabilized in the UbV3–PLpro complex (Fig. 5B). The measured K_D_ of UbV3-PLpro is 2.77 nM which is approximately 80-fold enhanced compared to UbV2 (A46F-K48E). Furthermore, IC_50_ of A46F-E64Y (34 nM) is 48-fold or 8-fold more efficient than A46F or E64Y, respectively. Both inhibition assays and binding affinities greatly suggest that the 3 positions (46, 48 and 64) cooperatively stabilize the interactions of Ub-PLpro hence UbV3 is a better inhibitor compared with UbV2 (Fig. 3C, 4). Altering surface charge (K48E and E64Y) or hydrophobicity (A46F and E64Y) on the small 76-residues Ub may lead to issues of structural stability. We thus employed circular dichroism (CD) spectroscopy to measure the far-UV CD spectrum of Ub and selected 9 UbVs at 25 and 80 °C to validate the thermostability of UbVs. The stability prediction also agrees with our experiments of UbV3 yielding high thermostability than single (A46F, K48E, E64Y) and double-point (UbV2) mutants (Fig. S9).

### Ub C-terminal mutations R74N-G75S further stabilize the PPIs

We next altered two residues, R75 and G75 of the Ub-tail region from our non-covalent network, which also serve a role in preventing cells from utilizing the UbVs. New sets of 74 and 75 mutations were added to our dual UbVs (Table 1), and the 4-point mutation A46F-K48E-R74N-G75S (termed UbV4) yielded a reduced IC_50_ of 110 nM. Similar with the double mutants, replacement of K48E by K48L/K48S/K48I undoubtedly results in the same inhibitory effects (IC_50_: 110-290 nM, Table S1). Therefore, we further included E64Y to UbV4 to use all 3 highly correlated regions and the designed 5-point mutant UbV5, resulting in an IC_50_ of 9.7 nM (5,500-fold of wtUb) and K_D_ of 1.5 nM (Table 1).

The characterized equilibrium constant K_D_ of UbV3 and UbV5 are approximately 20-fold enhanced compared to previously phage-display screened UbVs ME.2 and ME.4 (Table S1)^33^. In addition to impressive functional inhibition, UbV3 and UbV5 bound highly specifically to MERS PLpro (Fig. S10), which is also an essential attribute for a good inhibitor. Our UbV3 and UbV5 exhibit great inhibition equivalent to the reported variants for MERS PLpro^33^. In contrast to the preserved thermostability of UbVs designed in this study, the reported variants ME.2 and ME.4 with 15 mutated residues revealed denatured states at 80 °C^34^, which is significantly less thermostable than our UbVs (Fig S9). Similarly, T_m_ values of the 9-point mutant U7Ub25.2540 for USP7 is 64 °C lower than wtUb^34^. Mutating too many residues in Ub easily brings stability issues. Our work demonstrates the advantage and efficacy of modifying only a few residues of a protein template for increasing PPI.

In this study, we established a novel approach to computationally select key residues involving in the PPI based on the dihedral angle networks. We presented that the mapped dihedral angle networks are useful to identify critical interactions between two proteins. The identified residues were selected and mutated to alter the strengths of PPI. Using Ub-PLpro as the model system, we demonstrated that modifying a minimum of 2 to 3 residues within the correlation network in the Ub–PLpro interface can successfully enhance PPI and achieve 250-to 3,500-fold reduction of MERS PLpro activity. A combination of 5 mutated residues selected from the hydrophobic core, zinc finger region and the Ub-tail resulted in a 5,500-fold (IC_50_ = 9.7 nM) reduction in PLpro function and a 27,500-fold enhancement in UbV–PLpro complex affinity. Our design platform computationally examines a correlation network of protein sidechains and local pair-wise forces to efficiently design UbVs for experiments. The greatly correlated IC_50_ and K_D_ values (R^2^=0.94) imply that our experimental IC_50_ can be used to indirectly estimate K_D_ of UbV-PLpro. Integrating experimental measurements and further structural analysis using MD simulations and post-analysis iteratively re-informs new design.

The strategy is transformative for identifying key mutation sites and specific residues to guide rational design with high efficiency for many disease-linked DUBs and Ub-bound proteins^31,32^, including USP4^43^, USP7^44^, USP11^45^, and PLpro of SARS-CoV-2^31^. In contrast to producing variant-specific antibodies/vaccines for diverse spike proteins, blocking functions of viral nonstructural proteins is an alternative therapeutic solution to fight against the COVID-19. Designed tight and specific UbV inhibitors for the PLpro-CoV2 can provide a prominent solution to retard viral replication and rescue the antiviral immune response simultaneously. The same strategy can be straightforwardly applied to other protein-protein interacting systems related to signaling and enzymes. As these cellular events require frequent reoccurrences, protein complexes in cell signaling and enzymatic reactions do not have perfectly optimized PPI as steady bound complexes.

Therefore, mutating residues in these regions can remarkably improve binding. Modifying residues in regions exhibited highly correlated motions within the interaction network is also generalized and can be applied for engineering tightly bound variants for various applications.

## Materials and Methods

### MD simulation Protocol

X-ray crystal structure of the MERS-CoV-PLpro-wtUb was obtained from the RCSB protein data bank. (PDBID: 4rf0)^26^. PLpro was extended with 1 residue at the N-terminus and wtUb was extended with 2 residues to remain consistent residue numbers when comparing to preexisitng UbVs such as ME.2^33^. PLpro residues contain a total of 319 residues and Ub contains 78 residues. All MD simulations were performed by using the AMBER 20 package with GPU acceleration.^46^ Force Field ff14sb^47^ was used on proteins. First, we minimized the hydrogen atoms, amino acid side chain, and the entire protein system for 500,1000,5000 steps respectively in a general born implicit solvent. All systems were then solvated with TIP3P water with the extension of 12Å from the solute edge. Two counter-ion CL^-^ were added to neutralize the charge of the system. The solvated system contains roughly 72,000 atoms. The water molecules were minimized for 100-ps followed by the minimization of the whole system for 100ps. Third, the solvated system was equilibrated under constant pressure and temperature (NPT ensemble) from 50K to 275K with 25K increments and 100-ps each, and finally at 298K for 500-ps.

Production runs were also performed in NPT ensemble at 298K using Langevin Thermostat with 2-fs time step. The first 50-ns of MD simulation are treated as equilibrium plus, and thus Force distribution analysis (FDA)^37,38^ and molecular mechanics Poisson-Boltzmann surface area (MM/PBSA) calculations are performed using the following 450-ns. The cutoff of nonbonding interaction which includes Van Der Wall and Electrostatic components was set to 12 Å. The particle Mesh Ewald Method was used to compute long-range electrostatic interaction.

It is possible that a large protein-protein system is stuck at a specific local minimum and leads to suboptimal results. We performed three independent 150 ns MD production simulations for each conformation and selected the lowest energy trajectories by using MM/PBSA energy calculation. The exterior dielectric constant was set to 15 to accommodate the polar protein residues at the protein-water interface. The trajectories with the lowest energy were extended to a 500 ns production run. Output trajectories were saved every 1 ps for further analysis.

### Side Chain Dihedral Correlation Network

T-analyst is used to calculate the side chain dihedral rotations of each amino acid residue and their pairwise correlations. Each side chain dihedral angle was recorded every 100ps through 500ns trajectories which produce 5000 different angles per dihedral selection. The pairwise correlation which is computed using Pearson correlation formula requires angle corrections at the discontinuity margin (±180° or 360°/0°) or it will cause an error when computing their correlation ^48^. Positive correlation between two side chains indicates two sides rotate similarly during the MD simulation. We constructed the pair-wise correlation matrix between the side chain of wtUb and PLpro. From the pairwise correlation matrix, we generate a correlation network that allows us to visualize the correlation between each residue. Specific side chain rotation can generate a chain effect and affect the rotation of the distal residues. A correlation cutoff of 0.3 was applied to eliminate low correlated residues.

### PPI-GAMD Simulation Protocol

Starting from the last frame of our MD simulation, we perform 5-ns of classical MD simulation following by the 5-ns PPI-GAMD equilibration to correctly obtain the boost parameters. (ntcmdprep = 500,000, ntcmd=2500,000, ntebprep = 500,000, and nteb=2500,000 steps) Production runs were performed in NPT ensemble at 350K using Langevin Thermostat with 2-fs time step. We applied both potential boost and dihedral boost (igamd=17) on Ub residues that are within 5Å of the PLpro residues at the contact interfaces. Of note, applying dual boost potentials on entire Ub structure can result in denaturing protein structure and leads to suboptimal results. The threshold energy of potential boost is set to the upper bound limit (iEP=2) and the threshold energy of dihedral boost is set to the lower bound limit (iED=1). The upper limit of standard deviation of dual boost potential are set to 10 kcal/mol. (sigma0P and sigma0D=10) The production run continues until we observed Ub dissociate from PLpro. We defined the dissociation by observing a sudden jump of Cα RMSD values. The production runs were repeated with three different random seeds and the longest dissociation time were reported.

### Protein expression and purification

Genes including SARS-CoV-2 PLpro, MERS-CoV PLpro, ISG15 and ubiquitin were synthesized by GenScript (NJ, USA). Ubiquitin variants were made by site-directed mutagenesis or directed amplification (C-terminal mutations). UbV genes ME.2 and ME.4 were synthesized and subcloned by Genomics (Genomics Inc., Taiwan). All genes are placed in the pRSFDuet-1 vector with a N-terminal hexahistidine tag (his-tag) and a TEV cleavage sequence. All plasmids are transformed into BL21 RIL cell line for protein production. For PLpro and ISG15-AVTRYVDC, *E. coli* grown in LB medium at 37 °C with OD_600_ of 0.6-0.8 was induced by 0.6 mM isopropyl β-D-1-thiogalactopyranoside (IPTG) overnight at 16 °C (MERS-PLpro and SARS-CoV-2 PLpro) or 25 °C (ISG15-AVTRYVDC). For all 26 UbVs, *E*.*coli* was grown in the autoinduction medium containing the base broth (25 mM Na_2_HPO_4_, 25 mM KH_2_PO_4_, pH 7.2, 85 mM NaCl, 0.5% yeast extract, 2% tryptone) and a sugar mix (15% v/v glycerol, 1.25% w/v glucose, 5% w/v lactose) at a 25 : 1 volume ratio. The *E. coli* cells were cultured at 37 °C for 24 hours whereas UbV proteins were automatically expressed once glucose was depleted.

*E. coli* cell pellet was spun and resuspended with Buffer A (25 mM Tris-HCl pH 7.6, 200 mM NaCl, 3 mM 2-Mercaptoethanol) with the addition of 1 mM PMSF for sonication. Cell lysate was further centrifuged where the supernatant was loaded toward Roche cOmplete nickel resin followed by wash and eluted by 300 mM imidazole. The his-tag in PLpro was removed by TEV protease and proteins were further purified by size exclusion chromatography (SEC) on an Akta FPLC (Cytiva). ISG15-AVTRYVDC was further crosslinked with fluorescein-5-Maleimide (Santa Cruz Biotechnology) at 4 °C for 1 hour. Excess fluorescein was removed by desalting columns. All proteins were flash-frozen in liquid nitrogen, aliquoted and stored at -80 °C.

### Inhibition assay using fluorescence polarization (FP)

To detect and characterize the inhibitory effects on PLpro by the ubiquitin variants, we used fluorescein-labeled ISG15-AVTRYVDC (denoted as ISG15* hereafter) where AVTRYVD is the N-terminal sequence of SARS-CoV-2 and C was introduced from crosslinking reaction. The values of fluorescence polarization (FP) of ISG15* and the cleaved AVTRYVDC* are approximately 120 and 20, respectively (Fig. S7) which provide a sensitive tool to unravel the activity of MERS-CoV PLpro. 40 μl samples in multiple wells composed of 2 μM ISG15*, 50 nM MERS-CoV PLpro (or SARS-CoV-2 PLpro) and a wide range of UbV concentration (i.e. 0.06 nM – 150 μM) in a 384-well plate were measured for 1800 seconds or longer. ISG15* alone was used as a control. Individual UbV concentration was triplicated for statistical errors.

The reaction rate constants (*k*_*obs*_) of ISG15* cleavages were obtained by curve fitting using one-phase decay (equation 1). The enzymatic activity is normalized by the ratios of the *k*_*obs*_ values with and without UbV. To get the IC_50_ value, the normalized UbV concentration-dependent enzymatic activities are fitted by the logistic non-linear regression model (equation 2).

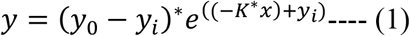

Where the y_0_ is the Y values when X (time) is zero, y_i_ is a plateau Y value at infinite times and *K* is the rate reaction constant.

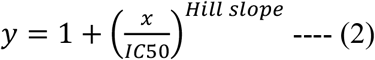

The Y values (the FP readouts) were normalized between 1.0 and 0 for the fitting where “Hill slope” was also included for fitting and the values were always around 1.0.

### Bio-Layer Interferometry (BLI)

BLI experiments are performed on the Octet RED 96 using anti-GST antibody biosensors for GST-tagged MERS-PLpro and UbV (or Ub) as analytes at 25 °C. The ligand and analytes are diluted into reaction buffer (25 mM Tris-HCl pH7.6, 150 mM NaCl, 0.1 mg/ml BSA, 0.01% tween-20). Steady-state response wavelength shifts of analytes in multiple concentrations are used to fit single-site binding system to get the dissociation constant (*K*_*D*_) using equation 3.

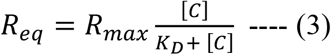

Where *R*_*eq*_ is the steady-state response shift of sensorgram curve, [C] is the analytes concentration, *R*_*max*_ is the maximal response and *K*_*D*_ is the dissociation constant. To determine *R*_*max*_ and *K*_*D*_ values, Levenberg–Marquardt algorithm is used to perform iterative non-linear least squares curve fitting.

The *k*_*on*_ and *k*_*off*_ values of UbV3 and UbV5 were globally fitted to the time-dependent response wavelength shifts in the association and dissociation well, respectively using the Octet Data Analysis software (Sartorius). *K*_*D*_ values of UbV3 and UbV5 are “*k*_*off*_ / *k*_*on*_”.

### Circular Dichroism (CD) Spectroscopy

CD measurements were performed on a Jasco spectropolarimeter J-815. Far-UV spectra were measured from 260 to 195 nm at 25 and 80 or 90 °C. All UbV samples containing 10 μM and diluted 25 mM Tris pH 7.6, 50 mM NaCl buffer were measured in a 1 mm quartz cell (Hellma GmbH). Melting temperature experiments were performed at 25 and 80 °C with an interval of 1 °C. The CD spectra were averaged from triplicates acquired at a scanning speed of 50 nm/min and a digital integration time of 1 second.

## Acknowledgments

This study was supported by the US National Institutes of Health (GM-109045), the US National Science Foundation (MCB-1350401 to C.C), and an Academia Sinica (Taipei, Taiwan) Career Development Award (AS-CDA-110-L03 to K.P.W.). We appreciate the technical and instrumental support from the Academia Sinica (AS) Biophysics Core Facility (AS-CFII-111-201) and the Biophysics Instrumentation Laboratory, Institute of Biological Chemistry, Academia Sinica.

## Author contributions

Conceptualization: KPW, CC

Methodology: TIH, YJH, WLL, KPW, CC

Formal Analysis: TIH, YJH

Data Curation: TIH, YJH, WLL

Supervision: KPW, CC

Writing—original draft: TIH, YJH, KPW, CC

## Code Availability

Source code for dihedral correlation network is available upon request.

## Data and materials availability

All data used in the main text or the supplementary materials are available on requests.

