## supporting information for "What Strengthens Protein-Protein Interactions: Analysis and Applications of Residue Correlation Networks"

**This PDF file includes:**

Supplementary Text
Figs. S1 to S10
Tables S1
References (1 to 5)

**Supplementary Text**

**MD Data Analysis**

**Optimize the Selection of Residues of Interest**

**1 Construct correlation network of sidechain dihedral angle for PLpro-Ub.wt.** To construct the correlation network, we analyze the 500-ns trajectories of wtUb in complex with PLpro. Each side chain dihedral angle was recorded every 100-ps through 500-ns trajectories which produce 5000 different angles per dihedral selection. We then calculate the pairwise correlation between each dihedral angles to construct the correlation matrix. With the correlation matrix, we construct the correlation network by using python library Networkx [1]. A correlation cutoff of 0.3 was applied to eliminate low correlated residues. We further eliminate the remaining residues that are outside the defined contact interfaces. (Figure S1)

**2 Define the Contact Interface and eliminate residues not at the contact interfaces.** FDA is used to select the contact interface between Ub.wt and PLpro. A cutoff of 10 (pN) was applied to disregard low interaction. We generate a heat map of pairwise forces between wtUb and PLpro. (Figure S2) From the heat map, four obvious interface regions were shown: Hydrophobic core, Alpha Helix region, Zn binding region, and BL2 region. Some highly correlated residues selected from the previous step are not located at the contact interface. Residues located outside the contact interface contribute little or no interaction to the binding affinity, and thus they are discarded.

**3 Consider multiple residues at the Zn binding region.** The Zn binding regions are highly fluctuated with high RMSF values and have been shown to crystallize in different conformations[2] The FDA also shows different patterns between different randomly seeded production runs. Of note, our dihedral correlation network selects S65 and T66, and the K63 is the conjugated side of PLpro. We believed that selecting E64 which is in the middle of S65 and K63 can optimize our result.

**4 Include ALA, GLY and PRO residues.** ALA, GLY and PRO residues do not have side chains, and thus will be neglected during dihedral angle selection. To avoid underestimating the possible mutation sites, we include all ALA, GLY and PRO residues within 5 Angstrom around our selected mutation sites of wtUb.

**5 Further select the remaining residues with FDA.** Residue of interest with strong attraction to the surrounding residues will not be selected for mutations. We focus on residues that are strongly repulsed or have very weak interactions to the surrounding. For instance, A46 has weak interaction with the surrounding residues, and K48 result in repulsive force with K204 of PLpro at the hydrophobic core. Therefore, A46 and K48 are ideal residues for mutation. On the other hands, FDA showed that residues 71 and 73 at the Ub-tail have strong attractive forces to the Alpha helix region, and thus mutation will not be applied in these three residues. (Figure S2) Finally, only 8 residues R42, A46, K48, E64, V70, R74 and G75 were selected for mutation.

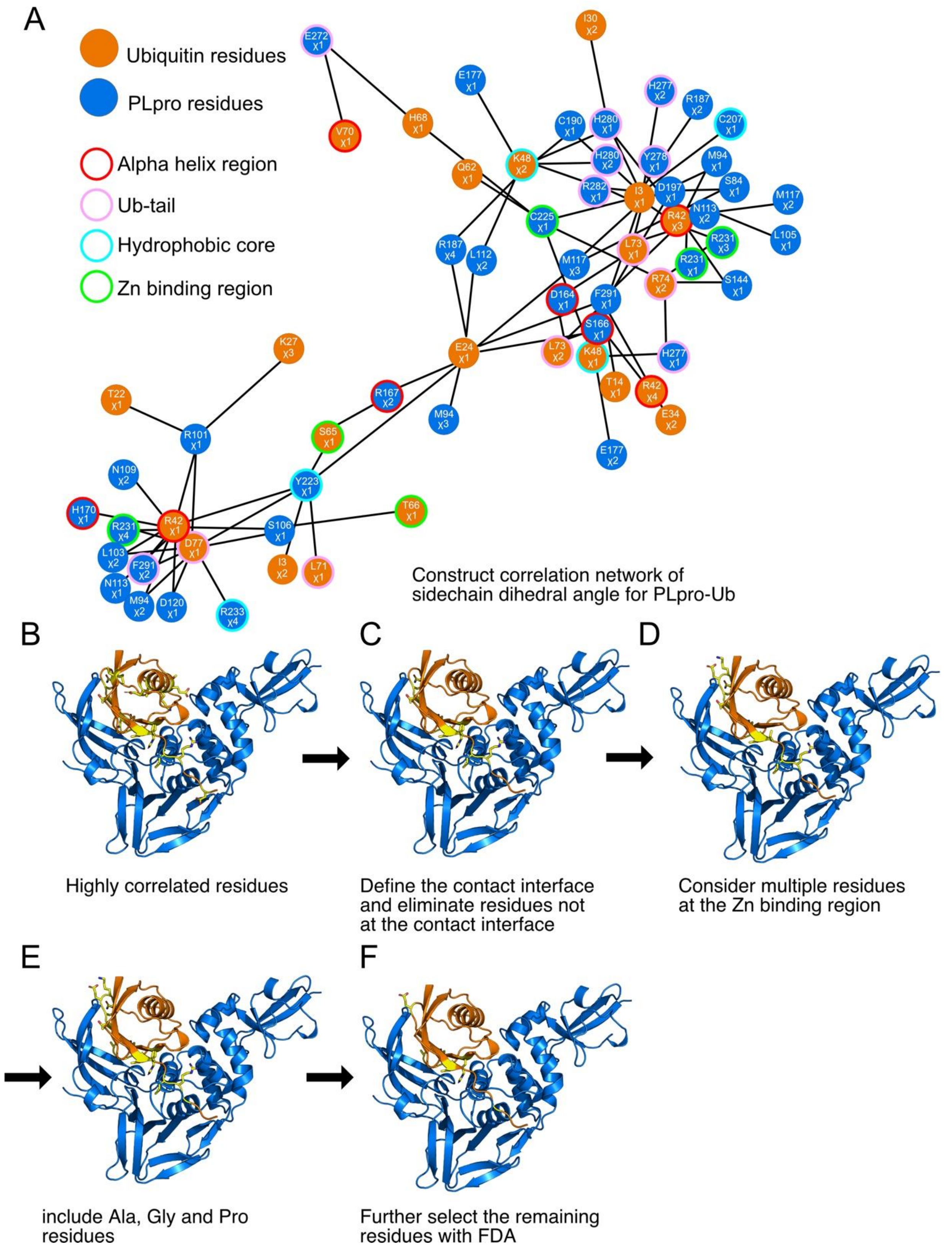
A
Ubiquitin residues
PLpro residues
Alpha helix region
Ub-tail
Hydrophobic core
Zn binding region
Construct correlation network of
sidechain dihedral angle for PLpro-Ub
B
Highly correlated residues
C
Define the contact interface
and eliminate residues not
at the contact interface
D
Consider multiple residues
at the Zn binding region
E
include Ala, Gly and Pro
residues
F
Further select the remaining
residues with FDA

**Figure S1. Optimized the Selection of Residues of Interest with side chain dihedral correlation network (A)** The side chain dihedral correlation network. The specific dihedral angles are indicated as $\chi$. **(B-F**) Detail steps of selecting the residues of interest. PLpro (marine), Ub (orange), and residues of interest (yellow).

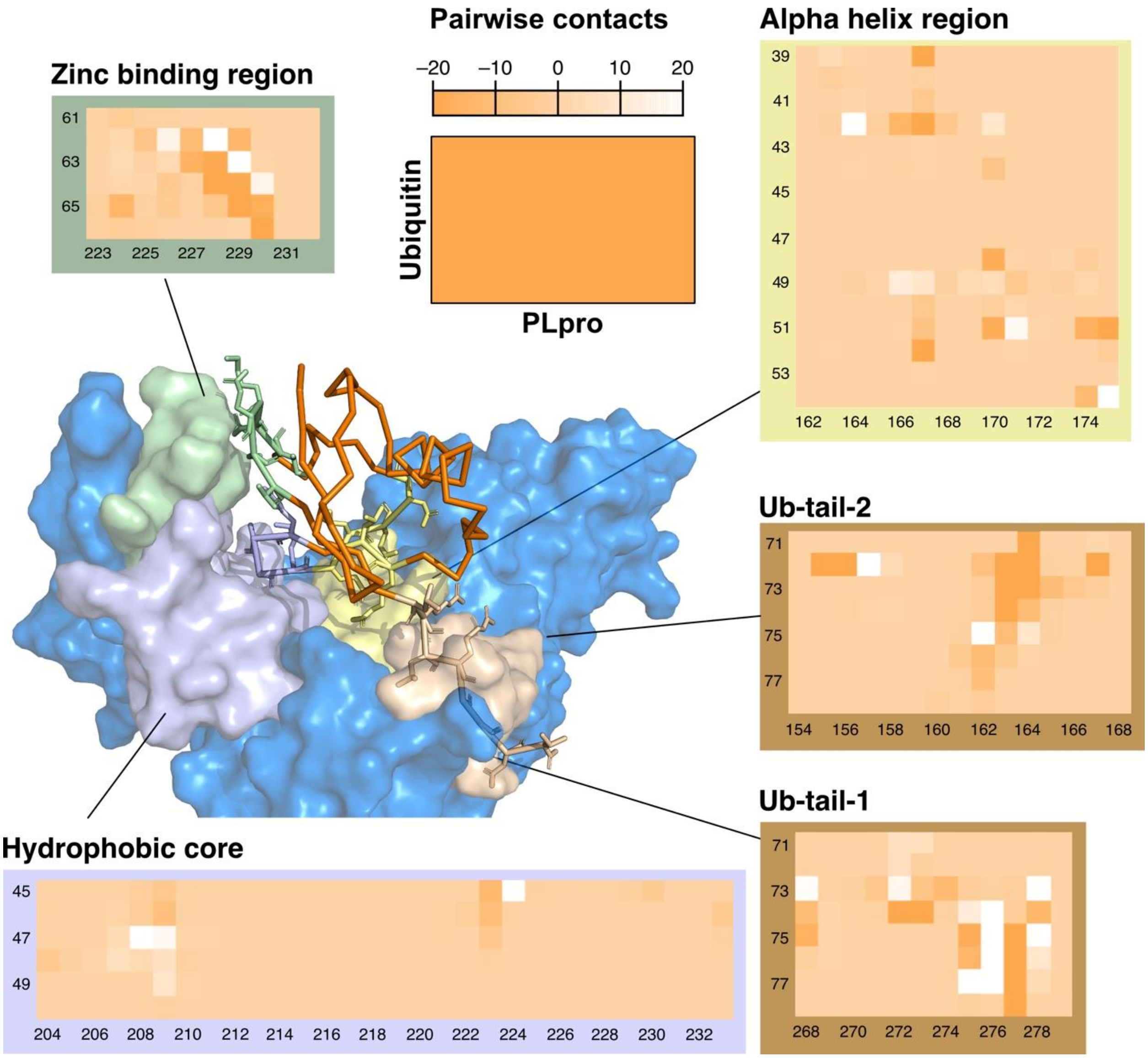


**Figure S2. Pair-wise forces distribution analysis (FDA) of contact regions between MERS-CoV PLpro and wtUb.** The FDA heatmap indicates if the pair-wise forces are attractive/negative or repulsive/positive. Four main contact regions are shown: Zn binding regions (palegreen), hydrophobic core (lightblue), alpha helix region (paleyellow), and Ub-tail (wheat).

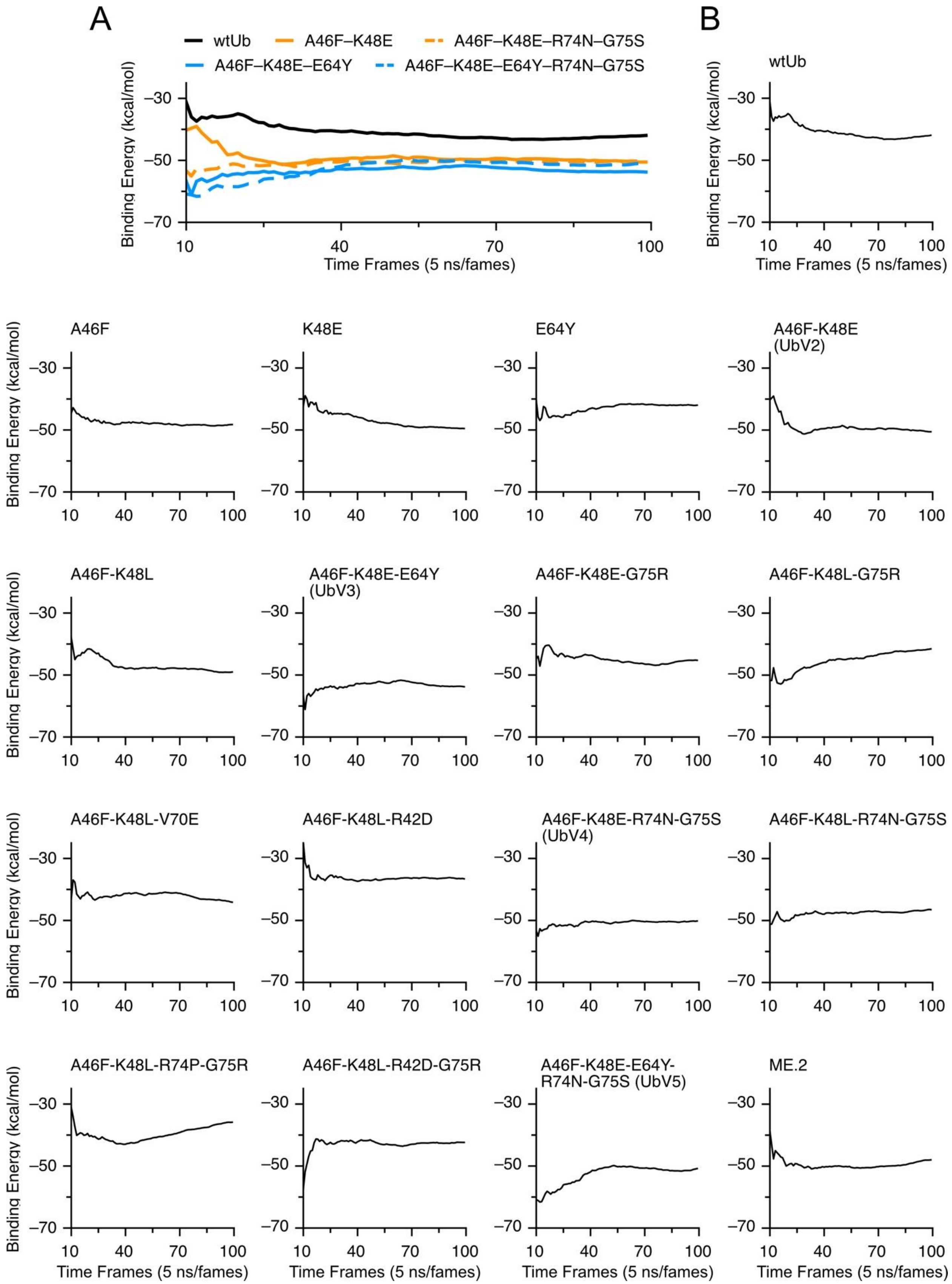

A
wtUb
A46F–K48E
A46F–K48E–R74N–G75S
A46F–K48E–E64Y
A46F–K48E–E64Y–R74N–G75S
Binding Energy (kcal/mol)
Time Frames (5 ns/fames)
B
wtUb
A46F
K48E
E64Y
A46F-K48E (UbV2)
A46F-K48L
A46F-K48E-E64Y (UbV3)
A46F-K48E-G75R
A46F-K48L-G75R
A46F-K48L-V70E
A46F-K48L-R42D
A46F-K48E-R74N-G75S (UbV4)
A46F-K48L-R74N-G75S
A46F-K48L-R74P-G75R
A46F-K48L-R42D-G75R
A46F-K48E-E64Y-R74N-G75S (UbV5)
ME.2

**Figure S3. MM/PBSA Block Analysis show convergences of binding energy of each UbVs.** To show convergence, we compute the mean binding energy at each time frames**. (A)** Comparing wtUb with our designed mutation, we observe that the binding energy of designed mutants are consistent lower than binding energy of wtUb throughout the MD simulation. **(B)** Block analysis of each UbVs shows convergence at the end of MD simulation.

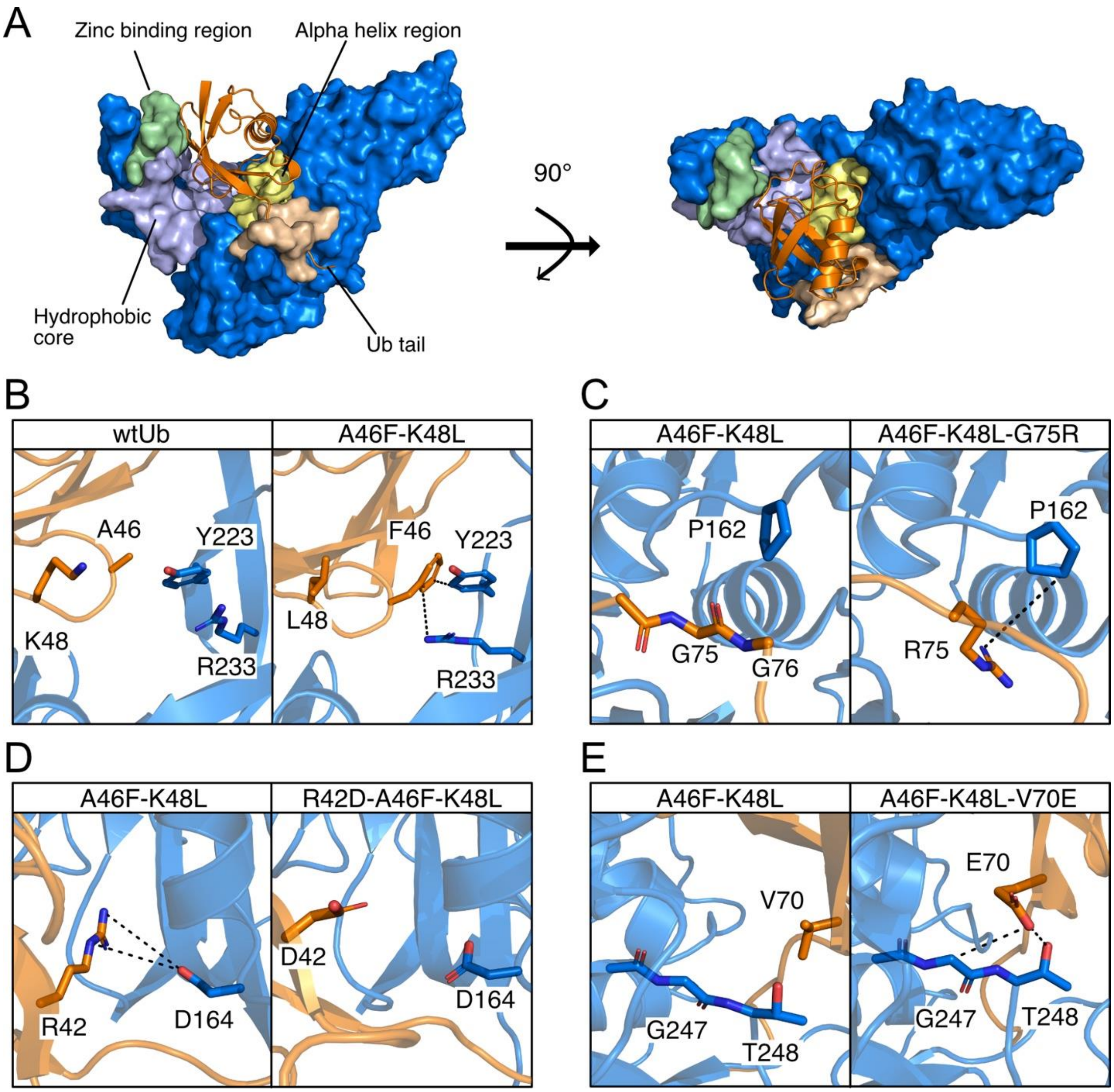


**Figure S4. Structural Analysis of each UbVs at different contact regions. (A)** Different perspective of MERS-CoV PLpro and Ub complex. **(B)** A46F mutation increase pi-pi with Y223 and pi-cation interaction with R233. **(C)** G75R mutation increase local vdW interaction due to the large carbon side chain. However, it results in a strong desolation penalty which is the energy require to remove the surrounding water molecules during binding. **(D)** R42D mutation disrupt the salt bridge between R42 and D164 which results in a significant decrease in binding affinity. **(E)** V70E mutation increase local attraction with G247 but stronger oxygen-oxygen repulsion was also observed which results in unfavorable binding affinity.

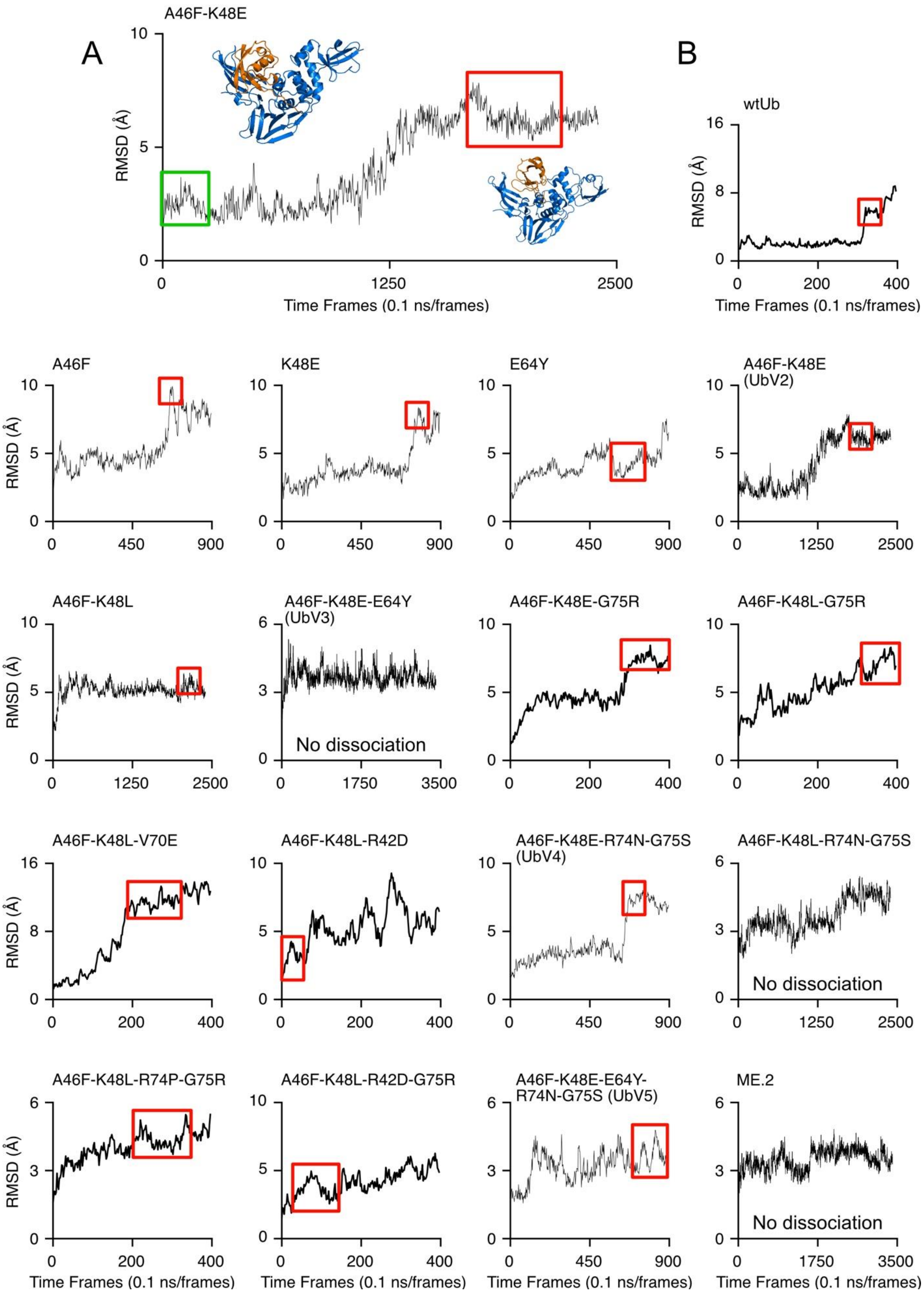
A
A46F-K48E
B
wtUb
A46F
K48E
E64Y
A46F-K48E (UbV2)
A46F-K48L
A46F-K48E-E64Y (UbV3)
No dissociation
A46F-K48E-G75R
A46F-K48L-G75R
A46F-K48L-V70E
A46F-K48L-R42D
A46F-K48E-R74N-G75S (UbV4)
A46F-K48L-R74N-G75S
No dissociation
A46F-K48L-R74P-G75R
A46F-K48L-R42D-G75R
A46F-K48E-E64Y-R74N-G75S (UbV5)
ME.2
No dissociation
RMSD (Å)
Time Frames (0.1 ns/frames)

**Figure S5. PPI-GAMD Dissociation Time of each UbVs.** PPI-GAMD allows us to observe the dissociation of UbVs from PLpro by applying boost potential on the contact interface of UbVs [3]. Root mean square deviation (RMSD) of the Ub-PLpro complex has been used to determine the bonded and unbonded states. Although a sudden jump of RMSD represents that Ub moves to a much more flexible state and is more likely to dissociate, we visualized the PPI-GAMD trajectories to capture the accurate time point where Ub dissociate from PLpro.

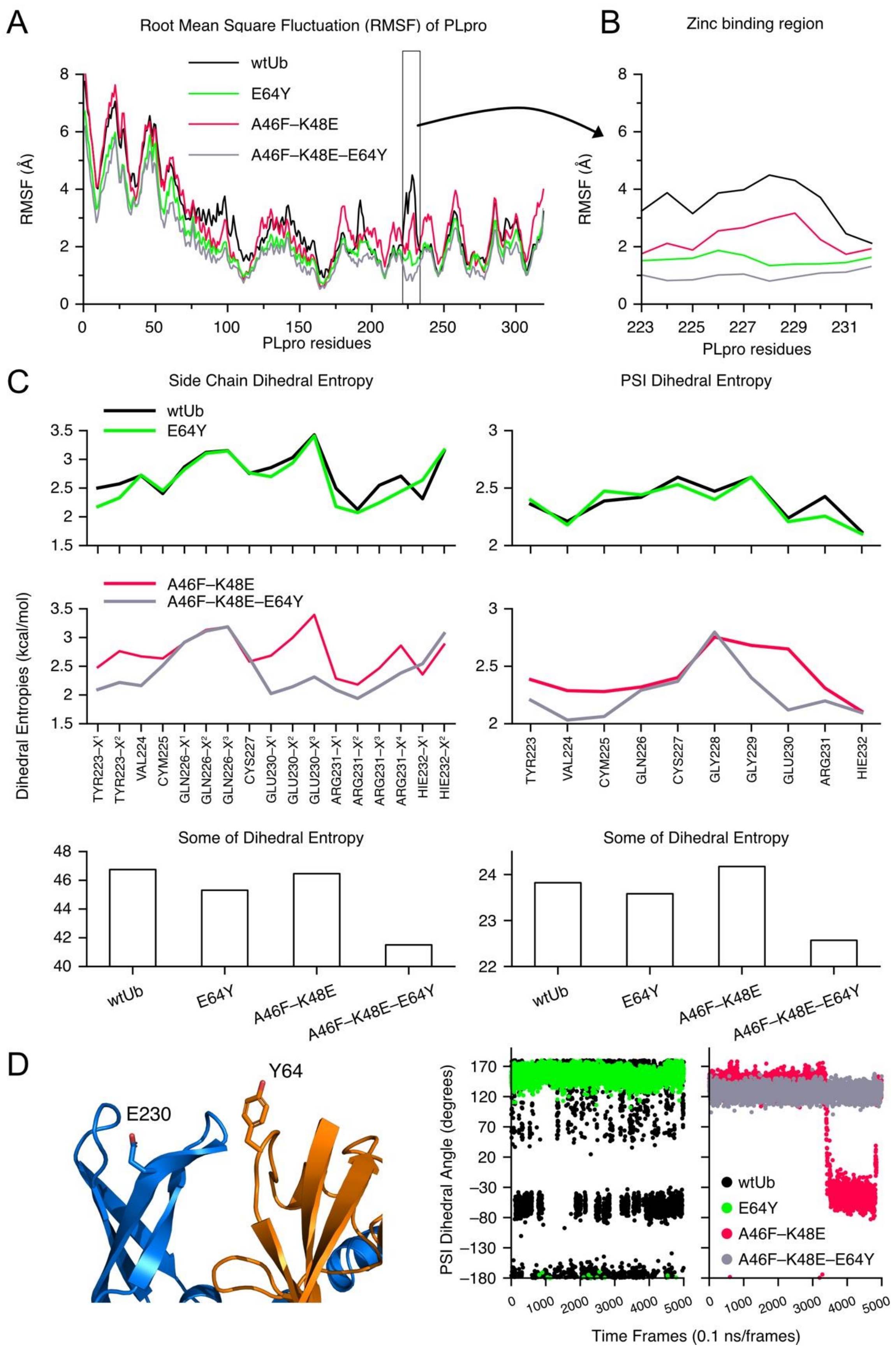
A
Root Mean Square Fluctuation (RMSF) of PLpro
wtUb
E64Y
A46F–K48E
A46F–K48E–E64Y
RMSF (Å)
PLpro residues
B
Zinc binding region
RMSF (Å)
PLpro residues
C
Side Chain Dihedral Entropy
PSI Dihedral Entropy
Dihedral Entropies (kcal/mol)
TYR223–X1
TYR223–X2
VAL224
CYM225
GLN226–X1
GLN226–X2
GLN226–X3
CYS227
GLU230–X1
GLU230–X2
GLU230–X3
ARG231–X1
ARG231–X2
ARG231–X3
ARG231–X4
HIE232–X1
HIE232–X2
TYR223
VAL224
CYM225
GLN226
CYS227
GLY228
GLY229
GLU230
ARG231
HIE232
Some of Dihedral Entropy
Some of Dihedral Entropy
wtUb
E64Y
A46F–K48E
A46F–K48E–E64Y
D
Y64
E230
PSI Dihedral Angle (degrees)
Time Frames (0.1 ns/frames)

**Figure S6. Quantifying the stability of E64Y mutations at the Zn binding regions. (A)** Cα RMSF of PLpro residues. **(B)** Zoom in at Zn binding region. E64Y mutation has lower RMSF values which indicate a reduction in local fluctuation. **(C)** Comparing the side chain and PSI dihedral entropies. Dihedral entropy is a configuration entropy using Gibbs entropies formula which uses the probability distribution of dihedral angle to further reveal local fluctuation.[4] The lower dihedral entropies indicate higher local stability. E64Y mutation reduced dihedral entropies and increase local stability. **(D)** PSI angle distribution of GLU230 which act as hinge. Hinge residues control the motion of flexible region. Without E64Y mutation, GLU230 of wtUb and A46F-K48E are able to adopt different dihedral rotations which result in higher flexibility at the Zn binding region.

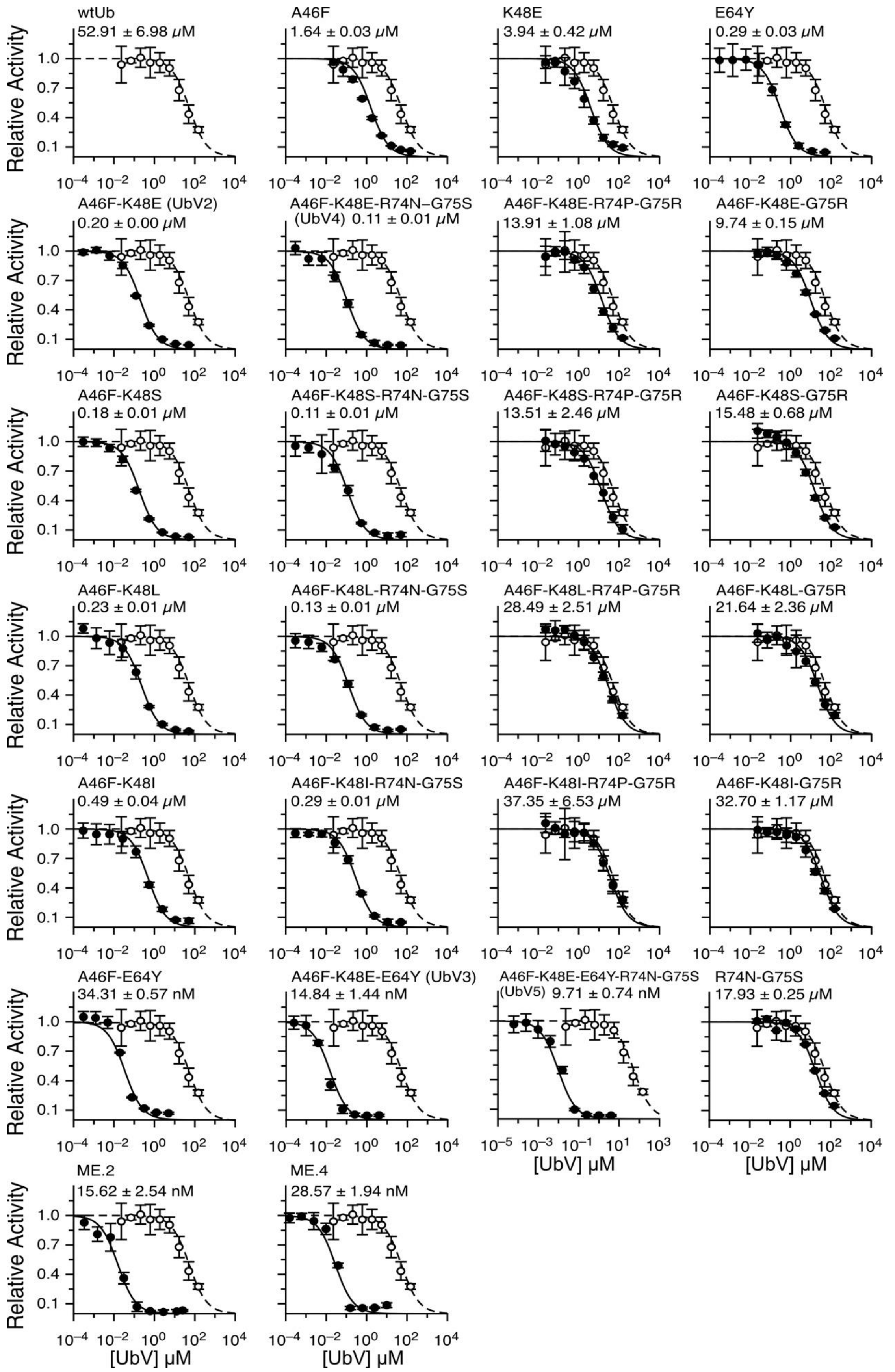
wtUb
52.91 ± 6.98 μM
A46F
1.64 ± 0.03 μM
K48E
3.94 ± 0.42 μM
E64Y
0.29 ± 0.03 μM
A46F-K48E (UbV2)
0.20 ± 0.00 μM
A46F-K48E-R74N–G75S
(UbV4) 0.11 ± 0.01 μM
A46F-K48E-R74P-G75R
13.91 ± 1.08 μM
A46F-K48E-G75R
9.74 ± 0.15 μM
A46F-K48S
0.18 ± 0.01 μM
A46F-K48S-R74N-G75S
0.11 ± 0.01 μM
A46F-K48S-R74P-G75R
13.51 ± 2.46 μM
A46F-K48S-G75R
15.48 ± 0.68 μM
A46F-K48L
0.23 ± 0.01 μM
A46F-K48L-R74N-G75S
0.13 ± 0.01 μM
A46F-K48L-R74P-G75R
28.49 ± 2.51 μM
A46F-K48L-G75R
21.64 ± 2.36 μM
A46F-K48I
0.49 ± 0.04 μM
A46F-K48I-R74N-G75S
0.29 ± 0.01 μM
A46F-K48I-R74P-G75R
37.35 ± 6.53 μM
A46F-K48I-G75R
32.70 ± 1.17 μM
A46F-E64Y
34.31 ± 0.57 nM
A46F-K48E-E64Y (UbV3)
14.84 ± 1.44 nM
A46F-K48E-E64Y-R74N-G75S
(UbV5) 9.71 ± 0.74 nM
R74N-G75S
17.93 ± 0.25 μM
ME.2
15.62 ± 2.54 nM
ME.4
28.57 ± 1.94 nM
Relative Activity
1.0
0.7
0.4
0.1
$10^{-4}$ $10^{-2}$ $10^{0}$ $10^{2}$ $10^{4}$
$10^{-5}$ $10^{-3}$ $10^{-1}$ $10^{1}$ $10^{3}$
[UbV] μM

**Figure S7. $IC_{50}$ plots of wtUb and 25 UbVs.** The wtUb as a weak competitive inhibitor is shown in open circles in each plot where UbVs are shown in black circles. The fitted $IC_{50}$ values of individual UbVs are provided on top of the corresponding fitting plots.

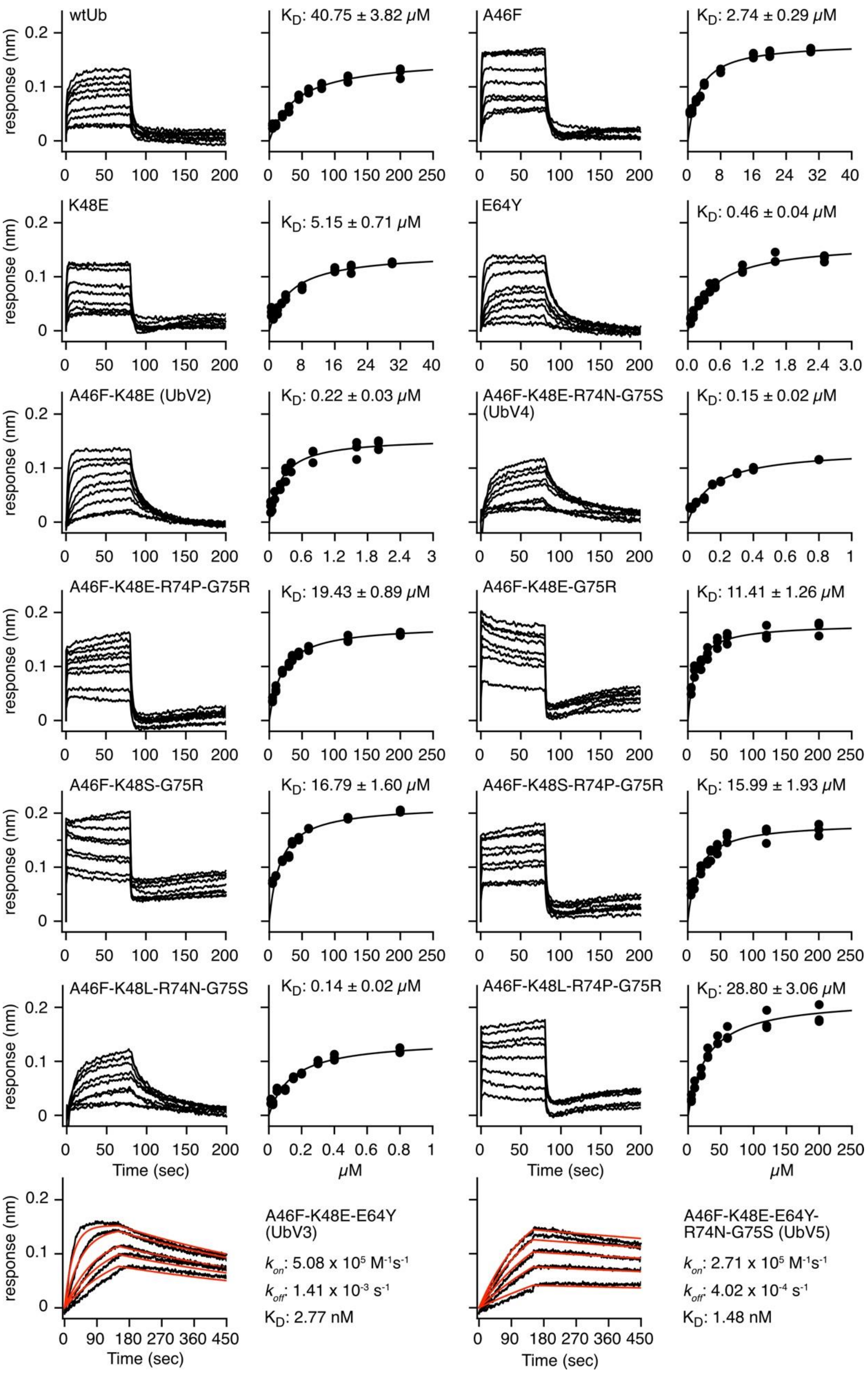
wtUb
K_D: 40.75 ± 3.82 μM
A46F
K_D: 2.74 ± 0.29 μM
K48E
K_D: 5.15 ± 0.71 μM
E64Y
K_D: 0.46 ± 0.04 μM
A46F-K48E (UbV2)
K_D: 0.22 ± 0.03 μM
A46F-K48E-R74N-G75S (UbV4)
K_D: 0.15 ± 0.02 μM
A46F-K48E-R74P-G75R
K_D: 19.43 ± 0.89 μM
A46F-K48E-G75R
K_D: 11.41 ± 1.26 μM
A46F-K48S-G75R
K_D: 16.79 ± 1.60 μM
A46F-K48S-R74P-G75R
K_D: 15.99 ± 1.93 μM
A46F-K48L-R74N-G75S
K_D: 0.14 ± 0.02 μM
A46F-K48L-R74P-G75R
K_D: 28.80 ± 3.06 μM
response (nm)
Time (sec)
μM
A46F-K48E-E64Y (UbV3)
k_on: 5.08 x 10^5 M^-1s^-1
k_off: 1.41 x 10^-3 s^-1
K_D: 2.77 nM
A46F-K48E-E64Y-R74N-G75S (UbV5)
k_on: 2.71 x 10^5 M^-1s^-1
k_off: 4.02 x 10^-4 s^-1
K_D: 1.48 nM

**Figure S8. Bio-layer interferometry (BLI) for the $K_D$ between MERS PLpro and UbVs.** The BLI sensorgrams of selected UbVs are shown along with the fitting curves using the steady-state responses. UbV3 and UbV5 are the two strongest binders and their $k_{on}$ and $k_{off}$ values were obtained by fitting the association and dissociation curves. $K_D$ was calculated by $k_{on}/k_{off}$.

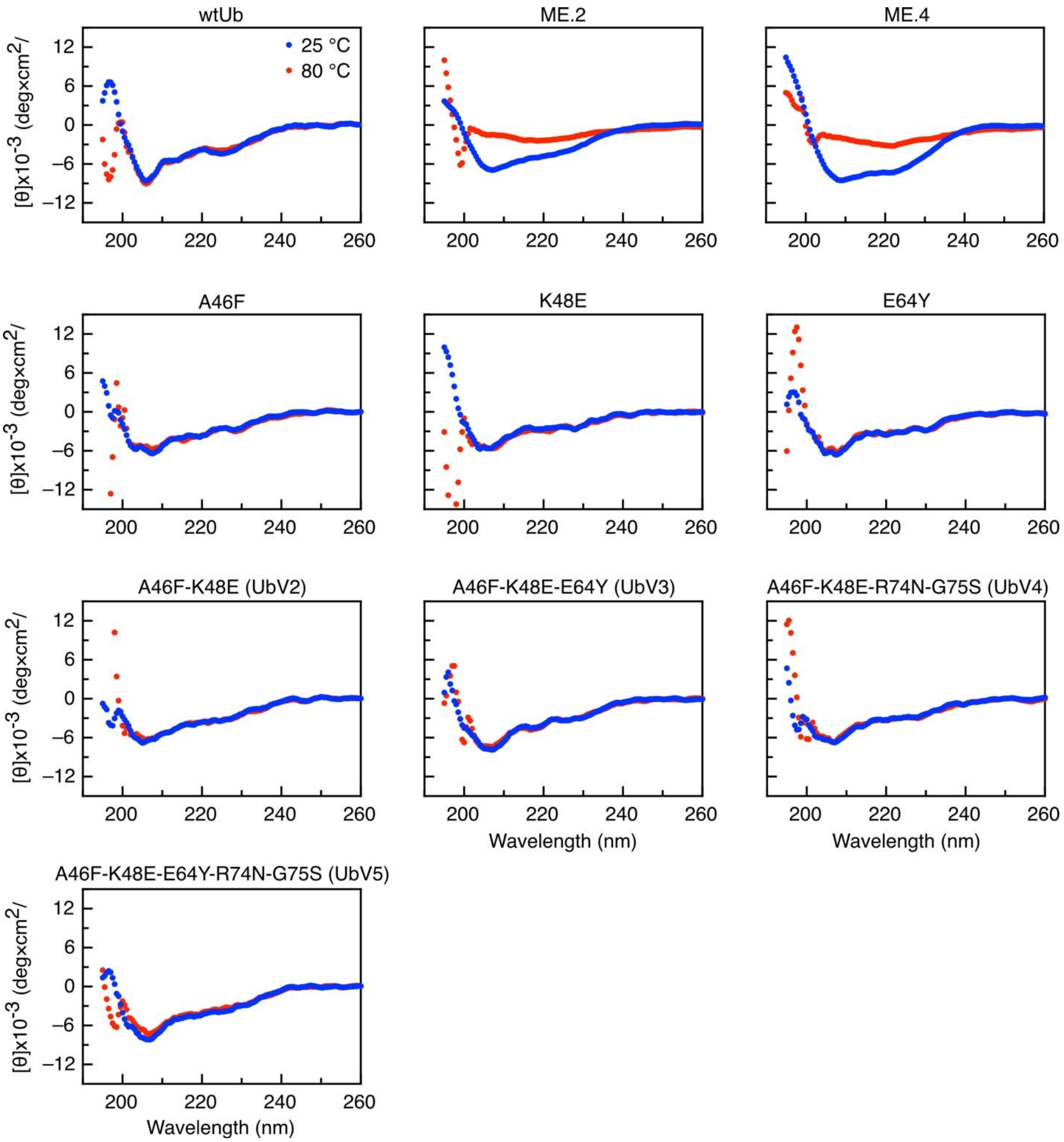


**Figure S9 Thermal stabilities of wtUb and UbVs.** The thermal stabilities of all proteins were monitored using far-UV circular dichroism (CD) spectroscopy at 25 and 80 ˚C. wtUb and designed UbVs in this study present identical CD curves at 25 and 80 ˚C suggesting the mutations do not interrupt the protein stability upon heating. On the contrary, previously selected UbVs ME.2 and ME.4[5] are not stable at 80 ˚C the secondary structural features in the CD curves substantially reduced.

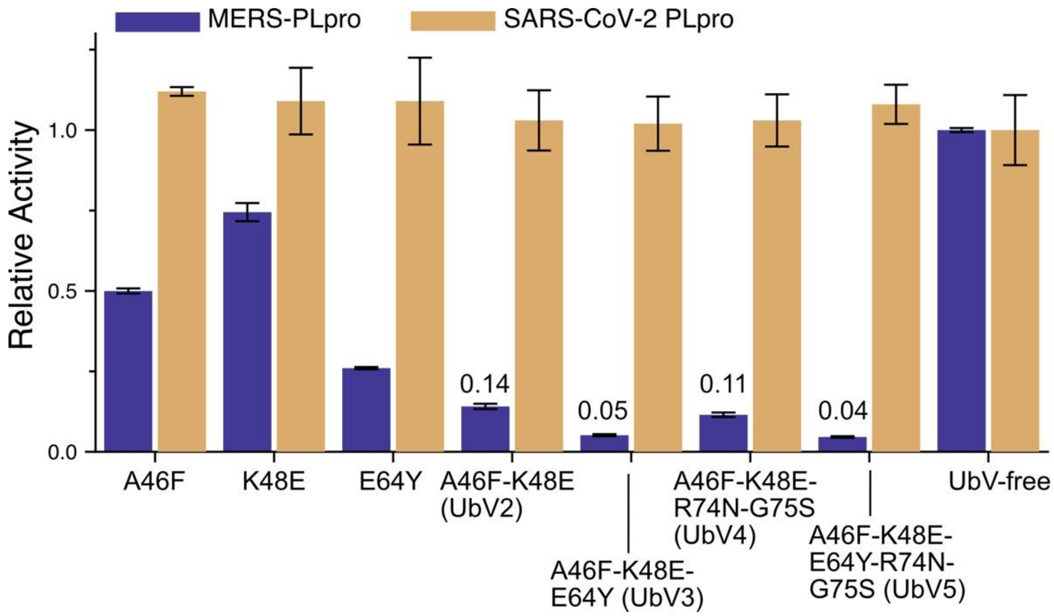


**Figure S10 The selectivity and potency of selected UbVs.** 7 UbVs were selected to validate the selectivity against MERS-CoV PLpro and SARS-CoV-2 PLpro. The PLpro activities were normalized to the UbV-free condition. The three single mutations present no interruptions to SARS-CoV-2 PLpro while the MERS-CoV PLpro activities were significantly reduced. Combinations of the 3 mutations and an addition of C-terminal mutations remain the same pattern. The UbV3 and UbV5 are substantially potent and specific to MERS-CoV PLpro. Remaining % activities of MERS PLpro were labeled for the multi-point mutation UbVs.

**Table S1. Computational and experimental evaluation of binding affinity between MERS-CoV-PLpro and all UbVs.**

| UbVs | VDW (kcal/mol) | EEL (kcal/mol) | Protein-protein (kcal/mol) | Protein-solvent (kcal/mol) | Binding Energy (kcal/mol) | Disso-ciation (ns) | $IC_{50}$ | $K_D$ |
|---|---|---|---|---|---|---|---|---|
| wtUb | -135.10 | -19.93 | -155.03 | 113.07 | -41.95 ± 2.56 | 32 | 52.91 ± 6.98 µM | 40.75 ± 3.82 µM |
| A46F | -142.62 | -14.9 | -157.53 | 109.28 | -48.25 ± 1.09 | 67.5 | 1.64 ± 0.03 µM | 2.74 ± 0.29 µM |
| K48E | -142.18 | -22.3 | -164.48 | 114.95 | -49.52 ± 2.56 | 75 | 3.94 ± 0.42 µM | 5.15 ± 0.71 µM |
| E64Y | -135.10 | -19.41 | -154.51 | 112.52 | -42.00 ± 1.49 | 68 | 0.29 ± 0.03 µM | 0.46 ± 0.04 µM |
| A46F-K48E **(UbV2)** | -144.5 | -22.43 | -166.93 | 116.39 | -50.54 ± 2.35 | 207 | 0.20 ± 0.00 µM | 0.22 ± 0.03 µM |
| A46F-K48L | -144.04 | -18.17 | -162.21 | 113.18 | -49.03 ± 2.33 | 200 | 0.23 ± 0.01 µM | N/A |
| A46F-K48S | -146.15 | -18.16 | -164.3 | 117.18 | -47.13 ± 2.84 | N/A | 0.18 ± 0.01 µM | N/A |
| A46F-K48I | NA | NA | NA | NA | NA | N/A | 0.49 ± 0.04 µM | N/A |
| K48E-V70E | -133.66 | -23.48 | -157.14 | 118.18 | -38.96 ± 1.96 | N/A | N/A | N/A |
| A46F-E64Y | N/A | N/A | N/A | N/A | N/A | N/A | 34.31 ± 0.57 nM | N/A |
| A46F-K48E-E64Y **(UbV3)** | -149.37 | -23.64 | -173.01 | 119.24 | -53.77 ± 1.37 | >250 | 14.84 ± 1.44 nM | 2.77 nM |
| A46F-K48E-G75R | -141.19 | -26.37 | -167.56 | 122.32 | -45.24 ± 1.48 | 40 | 9.74 ± 0.15 µM | 11.41 ± 1.26 µM |
| A46F-K48L-G75R | -147.82 | -19.01 | -166.82 | 125.29 | -41.53 ± 3.19 | 35 | 21.64 ± 2.36 µM | N/A |
| A46F-K48S-G75R | -135.9 | -21.43 | -157.33 | 114.23 | -43.10 ± 3.57 | NA | 15.48 ± 0.68 µM | 16.79 ± 1.60 µM |
| A46F-K48I-G75R | -146.26 | -22.02 | -168.27 | 122.58 | -45.69 ± 2.38 | NA | 32.70 ± 1.17 µM | N/A |
| A46F-K48L-V70E | -139.96 | -17.62 | -157.59 | 113.48 | -44.11 ± 1.13 | 27.5 | N/A | N/A |
| A46F-K48L-R42D | -130.75 | -18.36 | -149.11 | 112.52 | -36.58 ± 1.49 | 2.6 | N/A | N/A |
| **R74N-G75S** | N/A | N/A | N/A | N/A | N/A | N/A | 17.93 ± 0.25 µM | N/A |
| A46F-K48E-R74N-G75S **(UbV4)** | -143.08 | -20.73 | -163.81 | 118.61 | -50.18 ± 0.88 | 65 | 0.11 ± 0.01 µM | 0.15 ± 0.02 µM |

| UbVs | VDW (kcal/mol) | EEL (kcal/mol) | Protein-protein (kcal/mol) | Protein-solvent (kcal/mol) | Binding Energy (kcal/mol) | Disso-ciation (ns) | $IC_{50}$ | $K_D$ |
|---|---|---|---|---|---|---|---|---|
| A46F-K48E-R74P-G75R | N/A | N/A | N/A | N/A | N/A | N/A | 13.91 ± 1.08 µM | 19.43 ± 0.89 µM |
| A46F-K48L-R74N-G75S | -143.7 | -16.06 | -159.76 | 113.27 | -46.49 ± 0.98 | >250 | 0.13 ± 0.01 µM | 0.14 ± 0.02 µM |
| A46F-K48L-R74P-G75R | -128.48 | -19.62 | -148.1 | 112.24 | -35.86 ± 2.34 | 30 | 28.49 ± 2.51 µM | 28.80 ± 3.06 µM |
| A46F-K48L-R42D-G75R | -138.83 | -21.74 | -160.57 | 118.19 | -42.38 ± 2.09 | 8 | NA | NA |
| A46F-K48S-R74N-G75S | N/A | N/A | N/A | N/A | N/A | N/A | 0.11 ± 0.01 µM | N/A |
| A46F-K48S-R74P-G75R | N/A | N/A | N/A | N/A | N/A | N/A | 13.51 ± 2.46 µM | 15.99 ± 1.93 µM |
| A46F-K48I-R74N-G75S | N/A | N/A | N/A | N/A | N/A | N/A | 0.29 ± 0.01 µM | N/A |
| A46F-K48I-R74P-G75R | N/A | N/A | N/A | N/A | N/A | N/A | 37.35 ± 6.53 µM | N/A |
| A46F-K48E-E64Y-R74N-G75S **(UbV5)** | -149.04 | -20.38 | -169.42 | 113.62 | -50.81 ± 3.25 | 88 | 9.71 ± 0.74 nM | 1.48 nM |
| ME.2 | -148.37 | -18.96 | -167.32 | 119.35 | -47.98 ± 1.72 | >250 | 15.62 ± 2.54 nM | 53.2 ± 2.2 nM |
| ME.4 | N/A | N/A | N/A | N/A | N/A | N/A | 28.57 ± 1.94 nM | 35.9 ± 1.6 nM |
